# Effect of ORF7 of SARS-CoV-2 on the chemotaxis of monocytes and neutrophils in vitro

**DOI:** 10.1101/2021.09.13.460185

**Authors:** Gang Wang, Jun Guan, Guojun Li, Fengtian Wu, Qin Yang, Chunhong Huang, Junwei Shao, Lichen Xu, Zixuan Guo, Qihui Zhou, Haihong Zhu, Zhi Chen

## Abstract

Coronavirus disease 2019 (COVID-19) caused by the severe acute respiratory syndrome coronavirus 2 (SARS-CoV-2) is currently the most significant public health threats in worldwide. Patients with severe COVID-19 usually have pneumonia concomitant with local inflammation and sometimes a cytokine storm. Specific components of the SARS-CoV-2 virus trigger lung inflammation, and recruitment of immune cells to the lungs exacerbates this process, although much remains unknown about the pathogenesis of COVID-19. Our study of lung type II pneumocyte cells (A549) demonstrated that ORF7, an open reading frame (ORF) in the genome of SARS-CoV-2, induced the production of CCL2, a chemokine that promotes the chemotaxis of monocytes, and decreased the expression of IL-8, a chemokine that recruits neutrophils. A549 cells also had an increased level of IL-6. The results of our chemotaxis transwell assay suggested that ORF7 augmented monocyte infiltration and reduced the number of neutrophils. We conclude that the ORF7 of SARS-CoV-2 may have specific effects on the immunological changes in tissues after infection. These results suggest that the functions of other ORFs of SARS-CoV-2 should also be comprehensively examined.

## 1. Introduction

Coronavirus disease 2019 (COVID-19) is an infectious disease caused by the severe acute respiratory syndrome coronavirus 2 (SARS-CoV-2) that has become a worldwide pandemic [1-3]. The number of confirmed patients infected by SARS-CoV-2 continues to increase daily. As of Apr 2021, there were more than 0.14 billion SARS-CoV-2-infections and 3 million deaths from COVID-19 reported worldwide (https://coronavirus.jhu.edu/map.html). This global pandemic is still not under control, although there are encouraging trends in some regions. Other coronaviruses, such as MERS and SARS, had high transmissibility, but the epidemics were limited to certain regions and populations. Thus, SARS virus led to more than 8000 infected cases and 700 deaths in 26 countries and MERS led to about 2500 cases and 858 deaths in 27 countries [4-6]. In contrast, there has been an enormouse disease burden associated with SARS-CoV-2 infection. Numerous vaccines are currently available in many regions, and clinical trials have shown they are effective and safe [7, 8].

SARS-CoV-2 infects lung epithelial cells, type II alveolar (ATII) cells, by binding to the membrane-associated angiotensin-converting enzyme 2 (ACE2) on the cell surface [9-12]. Once inside the host cell, SARS-CoV-2 begins to produce viral RNA polymerase, which then replicates the complementary genomic RNA, making double-stranded RNA [13]. Subsequently, cells translate the structural and non-structural proteins (NSPs) of SARS-CoV-2 in the cytosol [13, 14]. The structural proteins include nucleocapsid (N), spike (S), membrane (M), and envelope (E) proteins, and the NSPs include 16 NSPs from the ORF1ab [15]. There are at least 10 open reading frames (ORFs) in the genome of SARS-CoV-2: ORF1ab, ORF2 (S protein), ORF3, ORF4 (E protein), ORF5 (M protein), ORF6, ORF7, ORF8, ORF9 (N protein), and ORF10 [15, 16]. Viral polymerase and all 16 NSPs are translated from the ORF1ab subgenome [16]. The S, N, E, and M proteins are all structural proteins, and the 16 NSPs function in replication and transcription of the viral genome [15].

Emerging evidence indicates that almost all of these ORF proteins have important roles in the lifecycle of SARS-CoV-2. As an RNA virus, SARS-CoV-2 infection stimulates the innate immunity of cells for RNA sensor proteins in the cytosol, such as retinoic acid inducible gene-I (RIG-I), melanoma differentiation-associated gene-5 (MDA-5), and toll-like receptors (TLR 3/7/8), which induce the expression of interferons (IFNs) [17]. ORF6, ORF8, and the N protein of SARS-CoV-2 inhibit these IFN-activated antiviral pathways, and this inhibits the IFN-stimulated response element (ISRE) [18]. Additionally, ORF3 upregulates markers of apoptosis in 293T, HepG2, and Vero E6 [19]. Although the functions of several ORFs are incompletely understood, all ORFs and NSPs have specific functions during the lifecycle of SARS-CoV-2.

The present *in vitro* study examined the function of ORF7 in SARS-CoV-2 by focusing on its regulation of numerous cytokines and chemokines (IL-6, TNF-a, IL-8, CXCL2, and CXCL7) that function in the chemotaxis of monocytes and neutrophils in vitro.

### 2. Materials and methods

### 2.1. Cell culture and vector construction of ORF7

A lung adenocarcinoma cell line (A549) was purchased from the Chinese Academy of Science (Shanghai, China) and cells were cultured in Dulbecco’s modified Eagle medium (DMEM; Hyclone, Logan, UT, USA) with 10% fetal bovine serum (FBS; Corning, NY, USA) and 1% penicillin/streptomycin in a humidified incubator with 5% CO_2_ at 37°C. For transfection, a lentiviral vector harboring FLAG-tagged ORF7 of SARS-CoV2 was constructed using a specific SARS-CoV-2 strain (Wuhan-Hu-1 strain, NC_045512). Transfection was performed and ORF7-expressing A549 cell was obtained. Control cells were transfected with control vector.

### 2.2. Quantitative real-time polymerase chain reaction (qRT-PCR)

RNA was extracted from cells using the Trizol reagent (TaKaRa, Dalian) and was then reverse transcribed into cDNA using the PrimeScript RT Master Mix (TaKaRa, Dalian). The expression of mRNAs (*IL-1a, IL-1b, IFN-α, IFN-β, IL-6, IL-8, CCL2*, and *TNF-α*) were quantified using qRT-PCR with the TB Green Master Mix (TaKaRa, Japan). Expression was normalized to *GAPDH*, and relative expression was calculated using the 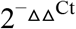 method (Primers are listed in Table 1). qRT-PCR was performed with the QuantStudio™ Dx system (ABI, Thermo, USA) using the following procedure: denaturation at 95°C for 5 min; 40 cycles of 95°C for 5 s and 60°C for 30 s; followed by a melting curve step of 95°C for 15 s and 60°C for 1 min, and a final increase to 95°C.

**Table 1.**
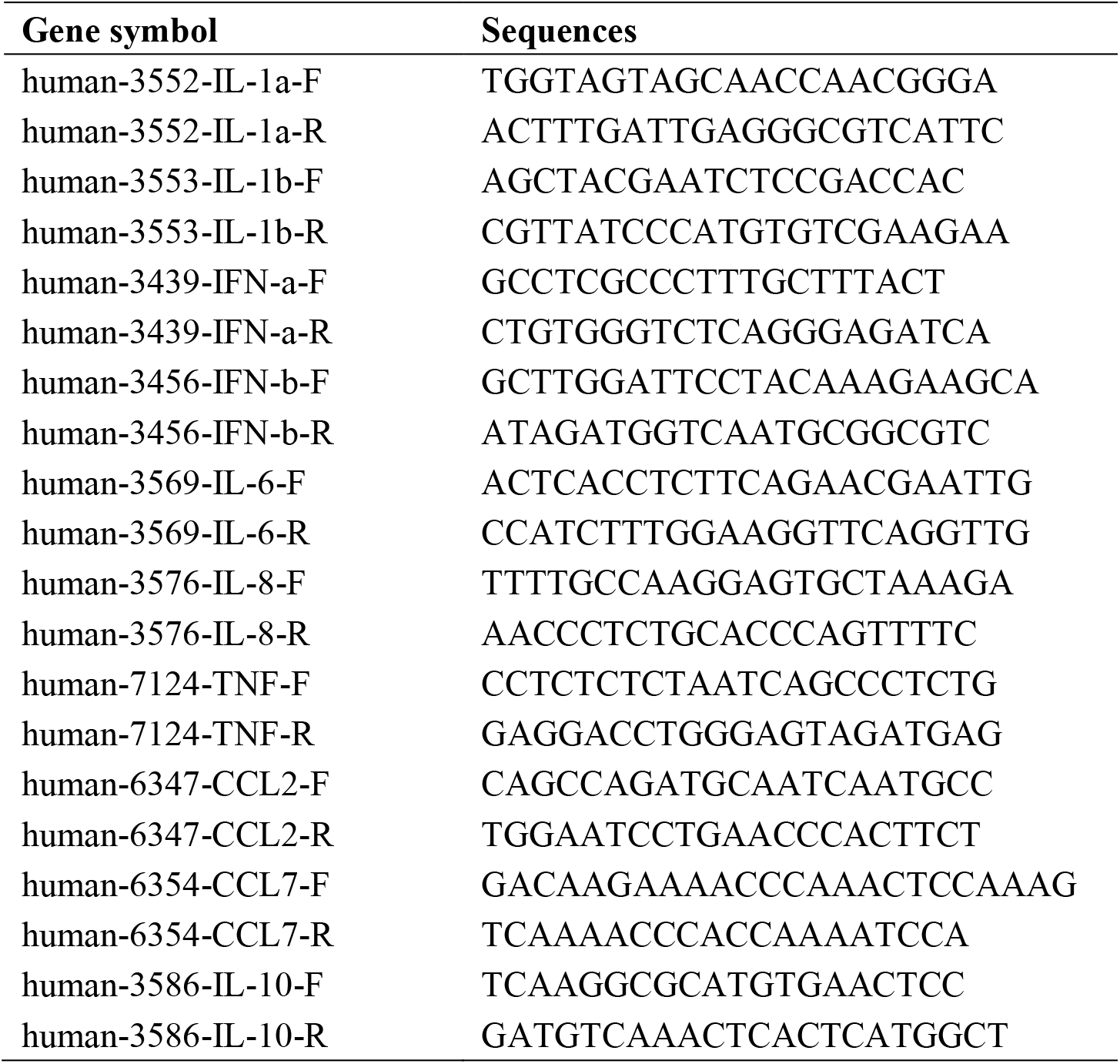
Primers for real time PCR.

### 2.3. Immunofluorescence

ORF7-expressing A549 cells were cultured in 6-well plates with slides. After 24 h, when the cells were adhered to coverslips, cells were fixed with 4% paraformaldehyde for 15 min, and permeabilized with 0.5% TritonX-100. After blocking for 30 min at room temperature using 3% BSA, the cells were incubated with a mouse anti-FLAG antibody (Sigma, USA) at 4°C overnight, and were then stained with a fluorescein isothiocyanate (FITC) conjugated goat anti-mouse antibody (Proteintech) at room temperature for 1 h. The nuclei were stained with DAPI (Abcam, ab104139) and the cellular distribution of ORF7 was observed using confocal microscopy.

### 2.4. Western blot analysis

Western blotting was conducted as previously described [20]. Briefly, cells were lysed with SDS sample buffer (1×), boiled for 10 min, separated using 4–20% SDS-PAGE (GenScript, USA), and then transferred onto a 0.22 μm polyvinylidene difluoride (PVDF) membrane (Millipore, USA). After blocking for 1 h at room temperature with 3% BSA, the membranes were incubated with diluted primary antibodies at 4°C overnight. The secondary antibodies were added at room temperature for 2 h. Protein bands were detected using the Clarity Western ECL Substrate (Bio-Rad, USA).

### 2.5. Isolation of human neutrophils and monocytes

Neutrophils were isolated from blood samples of healthy human donors using PolymorphPrep (Alere Technologies AS, Oslo, Norway) as previously described [21, 22]. Briefly, 5 mL of blood were layered onto 5 mL of PolymorphPrep and centrifuged for 35 min (500 *g*) at room temperature. The neutrophil layer was transferred to a new tube, washed with PBS, diluted by 50% with ddH_2_O, and then centrifuged for 10 min (400 *g*), followed by red blood cell lysis in a lysis buffer (Solarbio, China).

Peripheral blood mononuclear cells (PBMCs) were isolated from blood samples of healthy donors using a Ficoll density gradient [23]. Then, CD14 microbeads (Miltenyi Biotec, Germany) were used for the positive selection of human monocytes from these cells [24]. The purity of the CD14^+^ cells was evaluated using an APC-conjugated anti-human CD14 antibody (eBioscience, CatNo: 17-0149-42) with flow cytometry. The Institutional Ethics Committee of the First Affiliated Hospital of Zhejiang University approved this study.

### 2.6. Transwell assay

Transwell assays were conducted using a 12 mm transwell with a 3.0-μm pore polycarbonate membrane insert (Corning, USA, CatNo: CLS3402) [25]. A549 cells that were transfected with lentiviruses were seeded in the lower chamber and cultured for 24 h in DMEM containing 10% FBS. Human-derived monocytes and neutrophils in serum-free DMEM were added to the upper chamber. After incubation for 1 h, cells in the reverse side of the upper chamber were fixed with 4% paraformaldehyde and then stained in crystal violet for observation with a microscope. Cells in the lower chamber were collected and counted by flow cytometry, and counting beads were used for quantitation of different samples [26].

### 2.7. Enzyme-linked immunosorbent assay (ELISA)

ORF7-expressing A549 cells and control cells were plated into 6-well plates. The supernatant was collected and used for measurement of CCL2 (MultiSciences, CatNo: 70-EK187-96), CCL7 (cloud-clone corp, CatNo: SEA089Hu), and IL-6 (MultiSciences, CatNo: 70-EK206/3-96) using ELISA kits according to each manufacturer’s instructions.

### 2.8. Statistical analysis

The significance of differences was determined using Student’s *t*-test with GraphPad Prism version 7.0 (GraphPad Software, CA). A P value below 0.05 was considered significant.

## 3. Results

### 3.1. Construction of ORF-7-expressing A549 cells

ACE2 occurs on the surface of pneumocytes and binds to SARS-CoV-2 during infections [27]. We therefore first examined the expression of ACE2 in A549 cells, a type II pneumocyte cell line [28]. The western blotting and quantitative PCR results confirmed that these cells expressed ACE2 (Fig. 1A and B). We then used the sequence of a SARS-CoV-2 isolate (Wuhan-Hu-1, NC_045512.2) to construct a lentiviral vector that expressed a FLAG-tagged ORF7 subgenomic sequence (Lenti-ORF7-FLAG), and transfected A549 cells with Lenti-ORF7-FLAG to establish an ORF7-expressing cells. We confirmed the expression and the cellular distribution of ORF7 using western blotting and immunofluorescence. The results indicated expression of ORF7 (Fig. 1C) and that this protein was present in the cytosol (Fig. 1D). These data thus demonstrated the successful establishment of ORF7-expressing A549 cells that could be used for further studies of the function of ORF7.

**Figure 1.**
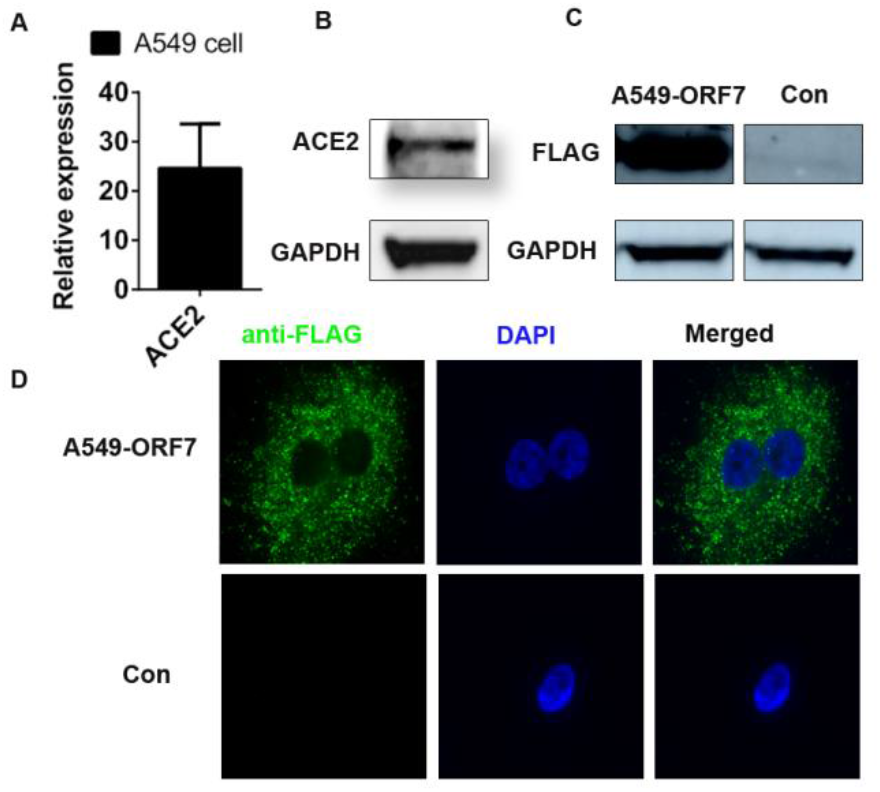
Expression of ACE2 and distribution of ORF7 in A549 cells. **A**, Expression of *ACE2* in A549 cells (qRT-PCR). **B**, Expression of ACE2 in A549 cells (western blotting). **C**, FLAG-ORF7 expression in A549-ORF7 cells (western blotting). **D**, Cellular distribution of ORF7 in A549-ORF7 cells (immunofluorescence microscopy).

### 3.2. ORF7 alters the expression of cytokines and chemokines in A549 cells

Innate immune cells, such as monocytes, macrophages, and neutrophils, are the first cells to respond when there is an infection in the lungs [29]. In particular, infection of pneumocytes leads to massive infiltration of monocytes into lung tissues [30], although there appears to be limited infiltration of neutrophils during SARS-CoV-2 infection [31-33]. During the early stage of viral infection, cytokines (IL-1, IL-2, IL-8, IL-10, CCL2, CCL7, IFN-α, IFN-β, and TNF-α) have essential functions in the recruitment of immune cells, defense against the infection, and promotion of inflammation [34]. We therefore used qPCR to determine the expression of cytokines and chemokines in ORF7-expressing A549 cells compared to control cells (Fig. 2A). The results demonstrated that *IL-6, CCL2*, and *IFN-β* had higher expression in A549-ORF7 cells (all P < 0.01), *IL-1α, IL-8*, and *TNFα* had lower expression in A549-ORF7 cells (all P < 0.01), the two groups had no differences in the levels of *IL-1-β* and *IFN-α* (both P > 0.05), and *CCL7* and *IL-10* were undetectable.

**Figure 2.**
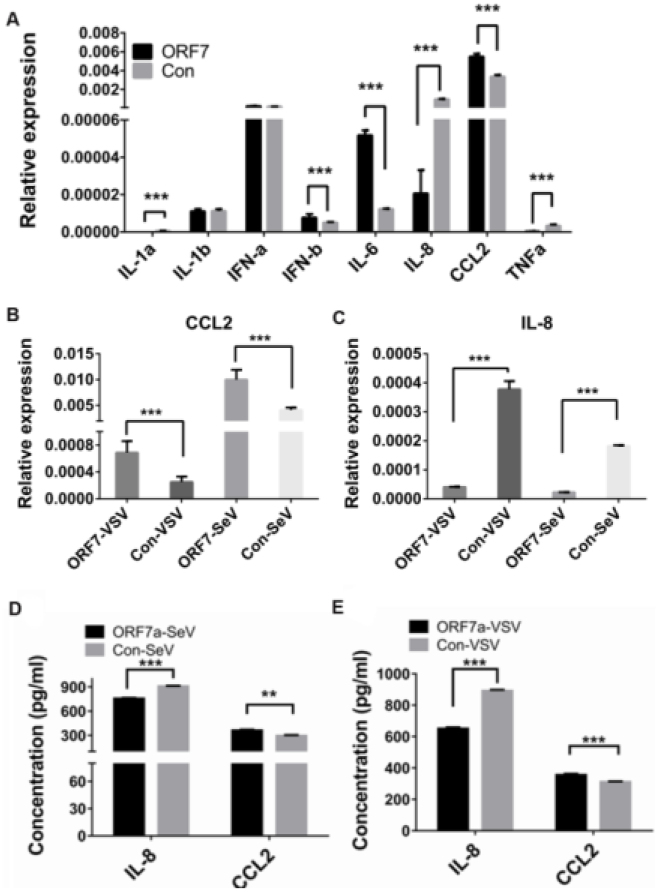
Effect of ORF7 on the expression of cytokines and chemokines in A549 cells. **A**, Relative to control cells, A549-ORF7 cells had increased expression of *IL-6, CCL2*, and *IFN-β* (all P < 0.01), decreased expression of *IL-1α, IL-8*, and *TNF-α* (all P < 0.01), but similar expression of *IL-1β* and *IFN-α* (both P > 0.05). *CCL7* and *IL-10* were undetectable (qRT-PCR). **B**, Activation of innate immunity by SeV/VSV infection increased the expression of *CCL2* and decreased the expression of *IL-8* in A549-ORF7 cells compared to control cells (both P < 0.05; qRT-PCR). **C**, A549-ORF7 cells had increased levels of CCL2 and decreased levels of IL-8 relative to control cells (both P < 0.01; ELISA).

CCL-2 (MCP-1) and IL-8 (CXCL8) function in the chemotaxis of monocytes and neutrophils, respectively [35, 36]. Sendai virus (SeV) and vesicular stomatitis virus (VSV) infections stimulate intracellular innate immunity, and can be used to model RNA virus infections [37]. We therefore used qPCR to measure *CCL2* and *IL-8* expression in A549-ORF7 and control cells following infection by these viruses (Fig. 2B). The results indicated that A549-ORF7 cells had greater expression of *CCL2* and decreased expression of *IL-8* compared to control cells (both P < 0.05). We also used ELISA to measure the levels of IL-8 and CCL2 in the supernatant of A549-ORF7 and control cells (Fig. 2C). These results confirmed that A549-ORF7 cells had increased expression of CCL2 and decreased expression of IL-8 with or without infection by SeV/VSV or the transduction of agonist, LMW and HMW RNAs (both P < 0.01).

### 3.3. ORF7 affects the chemotaxis of monocytes and neutrophils

We next examined the effects of ORF7 on the chemotaxis of monocytes and neutrophils. Flow cytometry provided identification of neutrophils as CD11b+ cells (Fig. 3A and B) and monocytes as CD14+ positive cells (Fig. 3C and D). Next, we implanted the A549-ORF7 and control cells in the lower chambers of 12-well plates, added monocytes and neutrophils in the upper chamber, and recorded chemotaxis after 6 h (monocytes) and 3 h (neutrophils) [25, 38]. The results indicated that monocytes had increased chemotaxis and neutrophils had reduced chemotaxis (Fig. 3 G–N).

**Figure 3.**
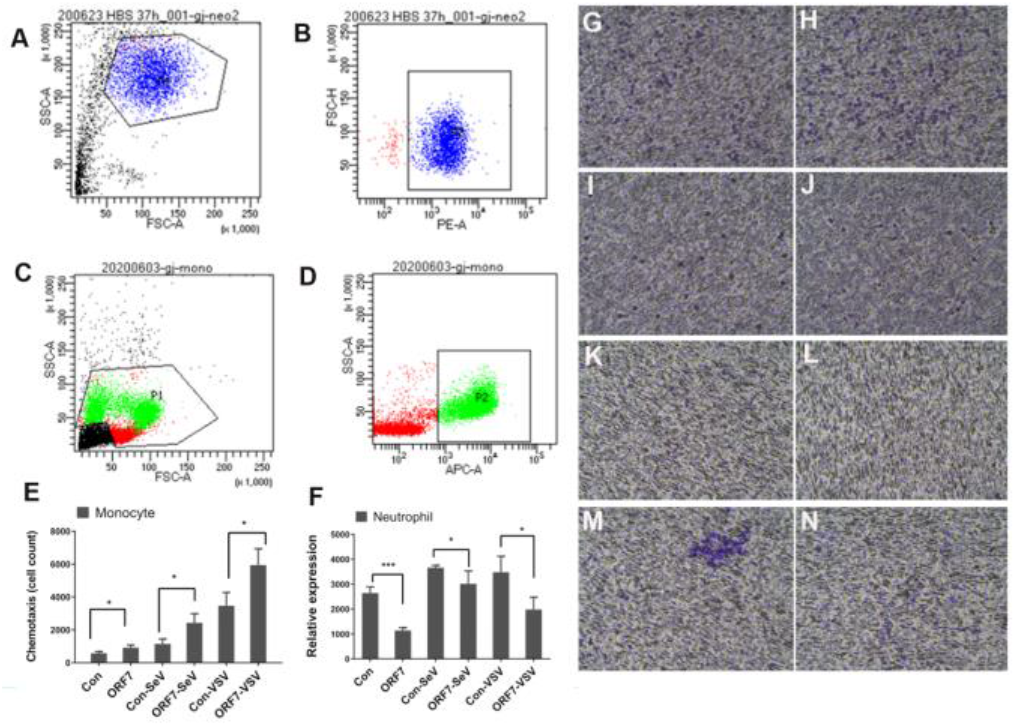
Effect of ORF7 on chemotaxis of neutrophils and monocytes. **A–D**, Neutrophils were identified as CD11b+ cells and monocytes as CD14+ positive cells (flow cytometry). **G–N**, Chemotaxis of monocytes and neutrophils (upper chamber) in response to A549-ORF cells (lower chamber) were examined by staining (transwell assay). **E–F**, Quantitation of chemotaxis results showing increased monocyte chemotaxis and decreased neutrophil chemotaxis (both P < 0.01).

During these experiments, we found numerous transmembrane cells, monocytes, and neutrophils on the lower wells with the A549-ORF7 cells. Thus, we used flow cytometry for the cell counting. The results indicated that more monocytes and fewer neutrophils migrated into the lower chamber with A549-ORF7 cells than with control A549 cells (all P < 0.01; Fig. 3E and F). These results indicated that ORF7 attracted monocytes and repelled neutrophils in vitro.

## 4. Discussion

Previous post-mortem examinations indicated that the lungs of COVID-19 patients, particularly the immune microenvironment, had significant alterations. These changes included alterations in T cells, B cells, macrophages, and neutrophils [39, 40]. Lymphocytes, T cells, and B cells were less abundant and scattered in the lungs of these patients [30, 32, 39, 41], but there was increased infiltration by monocytes, macrophages, and neutrophils [42]. After infection of the lungs, macrophages and neutrophils function as the first defense of the innate immune system, and these cells phagocytize pathogens and produce cytokines and chemokines that attract other immune cells [29]. The early recruiting of immune cells determines the local immune response, and can even cause more widespread inflammation, such as a cytokine storm. Our present work examined the influence of ORF7 of SARS-CoV-2 on innate immunity. Our major result is that expression of ORF7 in type II pneumocytes (A549 cells) increased the level of CCL2, decreased the level of IL-8, and increased the migration of primary monocytes but decreased the migration of neutrophils in vitro.

The ORF7 gene is located in a region of the genomes of the SARS-CoV-2, SARS-CoV-1, and MERS viruses that has a high frequency of mutations [1]. Our results indicated that ORF7 has a specific function in the immune response to coronavirus infection. Monocytes produce IL-1, IL-6, IL-18, IL-33, TNF-α, CCLs, and VEGF, and these molecules have critical roles in cytokine release and recruitment of other immune cells [43]. IL-6 and IL-1 are proinflammatory cytokines and the predominant inducers of the cytokine storm [43, 44]. Additionally, IL-6 can activate macrophages, which produce more cytokines and chemokines [43]. Neutrophils are also produced early in response to infection, and neutrophil chemotaxis in humans is usually mediated by factors such as IL-8, IL-1, TNF-α, and complement C5a [45]. Neutrophils, like macrophages, produce a range of cytokines (TNF-α, ILs, GCSF, MCSF, and GMCSF) and chemokines (IL-8, CXCL10, CXCL9, CCL2, CCL3, and CCL4) [43, 46]. Thus, neutrophils have direct anti-pathogen effects (phagocytosis) and indirect anti-pathogen effects (stimulation by cytokines). Macrophages and neutrophils thus play critical roles during the acute phase of pneumonia following viral infection.

Patients with COVID-19 have greater levels of peripheral monocytes than healthy controls [47, 48]. Recent evidence showed massive macrophage infiltration of the lungs of decreased COVID-19 patients, including intra-alveolar CD68+ macrophages [30, 31, 49]. The monocytes in these tissues (macrophages) may be at a stage of active proliferation in the lung alveolar spaces of these patients [50].

Previous research reported that the blood neutrophil count of SARS patients was associated with the severity of their pneumonia [51]. In contrast, other studies reported that the blood neutrophil count of COVID-19 patients was inversely associated with disease severity [47, 52, 53]. Although virus infections, such as influenza, induce neutrophil infiltration in the respiratory tract, the status of neutrophils in the lungs of COVID-19 patients appears paradoxical [30-33, 54, 55]. Several autopsy reports found no neutrophil infiltration in the lungs of COVID-19 patients, but there was neutrophil infiltration of the liver [31]. Some reports that found minor infiltrations of neutrophils in the lungs attributed this to secondary infections [33]. A report of 4 patients with COVID-19 found that only 1 patient had neutrophil infiltration of the lungs [55]. Another report of two cases found neutrophil infiltration in 1 patient’s pulmonary interstitium [30]. Other studies reported the presence of megakaryotes, dendritic cells, and natural killer cells in the lungs of deceased COVID-19 patients [33, 40].

The present *in vitro* study found that over-expression of the SARS-CoV-2 ORF7 in cultured type II alveolar cells (A549) up-regulated the expression of CCL2, a chemokine that functions in monocyte chemotaxis. This suggests that ORF7 may accelerate the progression of local inflammation after viral infection. Our *in vitro* studies also found that ORF7 downregulated the expression of IL-8. This suggests that ORF7 may block the migration of neutrophils. Greater neutrophil infiltration of the lungs could exacerbate the cytokine storm and worsen the patient’s condition due to lymphopenia [56].

There is evidence that a specific variant of SARS-CoV-2 which has mutation N501Y in the S protein and was first reported in London [57] is now widespread. Because this mutation is in the receptor-binding domain of the S protein, this variant likely has altered binding capacity to its ACE2 receptor. Even though this mutation was in the S protein, recent research reported the efficacy of neutralizing antibodies in mice [58]. It is important to consider that the mutation frequency of RNA viruses, such as coronaviruses and influenza viruses, are higher than that of DNA virus.

Our study indicated that ORF7 of SARS-CoV-2 had specific effects on the immunological changes. ORF7 may therefore contribute to the unique immunological profile of the lung tissues of patients with COVID-19, although the functions of other ORFs should also be examined. ORF7 can be the target for the development of anti-COVID-19 drugs in the future. The inflammation environment in the lung is one of the major risks for severe COVID-19 patients. In addition, ORF7 and other genes, including the structure and non-structure proteins, combinatorially involve in the inflammatory environment in the lung after the infection.

However, there are limitations in our work. Many other physical and pathological factors impact the inflammation status of the lungs of patients with SARS-CoV-2 infections. For example, the genomic RNA sensor system triggers the innate immune reaction, stimulates the translation of other proteins from the genome or subgenome of the virus, and may thereby increase lung injury. Consequently, appropriated animal models can be employed for the determination of ORF7 *in vivo* in the future. Additionally, the effect of ORF7 on the microenvironment of the lung should not be considered alone. Instead, there should be a systematic examination of the impact of all factors and their interactions on lung inflammation.

## Data Availability

There are no publicly archived datasets analysed in this study.

## Ethical Approval

Our study was approved by the Institutional Ethics Committee of the First Affiliated Hospital of Zhejiang University

## Consent

Written informed consent of the legal guardian was obtained.

## Author Contributions

Conceptualization, Z.C. and H.Z.; Methodology, G.W., J.G., F.W., C.H. J.S., L.X., Z.G., and Q.Z.; Formal Analysis, G.W. and J.G.; Data Curation, G.W. and J.G.; Writing Original Draft Preparation, G.W. and J.G.; Writing – Review & Editing, Z.C., H.Z. and G.L.; Supervision, Z.C. and H.Z.; Project Administration, Z.C.; Funding Acquisition, Z.C.

